# Genome Restructuring around Innate Immune Genes in Monocytes in Alcohol-associated Hepatitis

**DOI:** 10.1101/2024.08.07.607014

**Authors:** Adam Kim, Megan R. McMullen, Annette Bellar, David Streem, Jaividhya Dasarathy, Nicole Welch, Srinivasan Dasarathy

## Abstract

Many inflammatory genes in the immune system are clustered in the genome. The 3D genome architecture of these clustered genes likely plays a critical role in their regulation and alterations to this structure may contribute to diseases where inflammation is poorly controlled. Alcohol-associated hepatitis (AH) is a severe inflammatory disease that contributes significantly to morbidity in alcohol associated liver disease. Monocytes in AH are hyper-responsive to inflammatory stimuli and contribute significantly to inflammation. We performed high throughput chromatin conformation capture (Hi-C) technology on monocytes isolated from 4 AH patients and 4 healthy controls to better understand how genome structure is altered in AH. Most chromosomes from AH and healthy controls were significantly dissimilar from each other. Comparing AH to HC, many regions of the genome contained significant changes in contact frequency. While there were alterations throughout the genome, there were a number of hotspots containing a higher density of changes in structure. A few of these hotspots contained genes involved in innate immunity including the NK-gene receptor complex and the CXC-chemokines. Finally, we compare these results to scRNA-seq data from patients with AH challenged with LPS to predict how chromatin conformation impacts transcription of clustered immune genes. Together, these results reveal changes in the chromatin structure of monocytes from AH patients that perturb expression of highly clustered proinflammatory genes.

## Introduction

The three-dimensional (3D) architecture of chromatin in the nucleus is highly organized and contributes to the regulation of gene expression and cellular differentiation. Within the nucleus, each chromosome forms compartments that are distinct and separate from each other, called chromosome territories^1^. Within chromosome territories, the 3D architecture forms many distinct local features, like loops and topologically associating domains (TADs), as well as long-range interactions between enhancers, insulators, and promoters^2^. Cohesin and CCCTC-binding factor (CTCF) determine the structure and boundaries of TADs and these structural elements are conserved in many different cell types and species^3,4^. Loss of these proteins in most cell types disrupts TAD structure and chromatin loops although surprisingly gene expression is mostly unchanged^5–7^. But in myeloid cells, loss of CTCF greatly disrupts expression of innate immune genes after acute inflammatory stimuli, suggesting local chromatin structure has a significant role in certain context specific gene expression responses^8,9^.

While transcription factors, like nuclear factor-kappa B (NF-κB) and interferon-regulatory factors (IRF), bind and activate specific regulatory elements in the genome^10^, the 3D genome architecture, via TADs and chromatin loops, bring these elements into close proximity, and increase associations between enhancers and promoters of genes in similar activation states^11,12^. In certain contexts, transcription factor activation can then alter the 3D genomic landscape by increasing chromatin accessibility and contact frequency of certain regions. For example, in response to type 1 and type 2 interferons, loci in the genome containing families of interferon stimulated genes, increase chromatin contacts^13^.

While the 3D architecture plays a major role in cellular differentiation and gene regulation, less is known about how chromatin structure changes in disease^11,14^. A recent study, which performed high-throughput chromatin conformation capture technology (Hi-C) on primary monocytes from two patients with systemic lupus erythematosus (SLE) and two age- and sex-matched healthy controls, found few changes in chromatin structure between healthy and disease individuals^15^. This would suggest that Hi-C might not be a useful tool for predicting how monocytes transform in disease, and how genome architecture influences gene expression in primary cells. On the other hand, a few studies have found many differences in chromatin structure between primary monocytes and THP-1 cells, a monocytic cell line^15–17^. While these data suggest that 3D genome architecture may be a conserved feature of cellular differentiation and not a significant contributor to disease, more studies have to be conducted in different disease contexts to see how chromatin structure can influence monocyte gene expression. One limitation of the previous studies is that, due to the cost of these experiments, the sample size tends to be low, 1-2 patient samples, and data is often combined to increase resolution. As a result, statistical testing and disease specific comparisons have not been measured in most previous studies.

Alcohol-associated Hepatitis (AH) is a severe inflammatory disease characterized by extremely pro-inflammatory immune cells which infiltrate and damage the liver^18,19^. Monocytes in AH express significantly more pro-inflammatory genes in response to innate immune stimuli^20–22^. Many innate immune genes are clustered in the genome, like the CXC-chemokine cluster on Chromosome 4, the CC-chemokine cluster on Chromosome 17, and the NK-gene receptor complex on Chromosome 12, all of which have been implicated in AH. Single cell analysis revealed that these genes have highly coordinated expression patterns, suggesting the proximity of these genes influences their expression patterns^20^.

Here, we present the 3D genome architecture of primary monocytes isolated from four patients with severe AH, and four age-matched healthy controls using Hi-C technology. We hypothesized that hypersensitivity to bacterial lipopolysaccharides (LPS) and heightened pro-inflammatory responses in AH might be due to a significantly perturbed 3D genome architecture. Our results show that there are extensive changes in chromatin structure throughout the genome in AH. While changes in contact frequency occurred throughout every chromosome, there were a number of hotspots with a high density of changes, and in many cases, hotspots contained important families of genes involved in immunity and have been implicated in AH. Ultimately, these results suggest the 3D genome structure is significantly altered in AH, and these changes contribute to perturbed gene expression patterns, specifically pro-inflammatory genes.

## Results

### Hi-C reveals the 3D genome architecture of hyper-inflammatory monocytes from AH patients

To investigate changes in 3D genome architecture, we performed Hi-C on primary monocytes isolated from four AH patients (all males) and 4 age-matched healthy controls (3 males and 1 female, **Table 1, Supplemental Table 1**). Cryo-preserved PBMCs were thawed and CD14+ and CD16+ monocytes were isolated by negative selection. Hi-C was then performed using the Arima kit, and all 8 libraries were sequenced to an average depth of about 570 million reads per sample (**Supplemental Table 2**). Using Juicer, the reads were aligned to the genome and chromatin conformation was determined at a resolution of 100kb. As expected, most of the chromatin interactions observed were local and intra-chromosomal (**Figure 1, Supplemental Figure 1, Supplemental Table 2**). As a result, we focused on intra-chromosomal analyses and not inter-chromosomal interactions.

**Table 1:** Summary of Clinical data for patient samples used for Hi-C.

| Table 1: Summary of Clinical data for patient samples used for Hi-C |  |  |
| --- | --- | --- |
|  | Healthy Control<br>Lean, n=4 | Alcohol-associated<br>Hepatitis Lean, n=4 |
| Age (years) | 45 (31-57) | 46 (25-62) |
| Sex | 3M/1F | 4M |
| AST (U/L) | - | 115 (30-241) |
| ALT (U/L) | - | 44 (21-99) |
| Serum Albumin (g/dL) | - | 3.1 (3.0-3.3) |
| Serum Creatinine (mg/dL) | - | 1.81 (1.04-3.50) |
| Serum Bilirubin (mg/dL) | - | 21.5 (1.9-30.9) |
| HBA1c | - | 9.1 (7.2-10.5) |
| MELD Score | - | 26 (14-35) |

**Figure 1:**
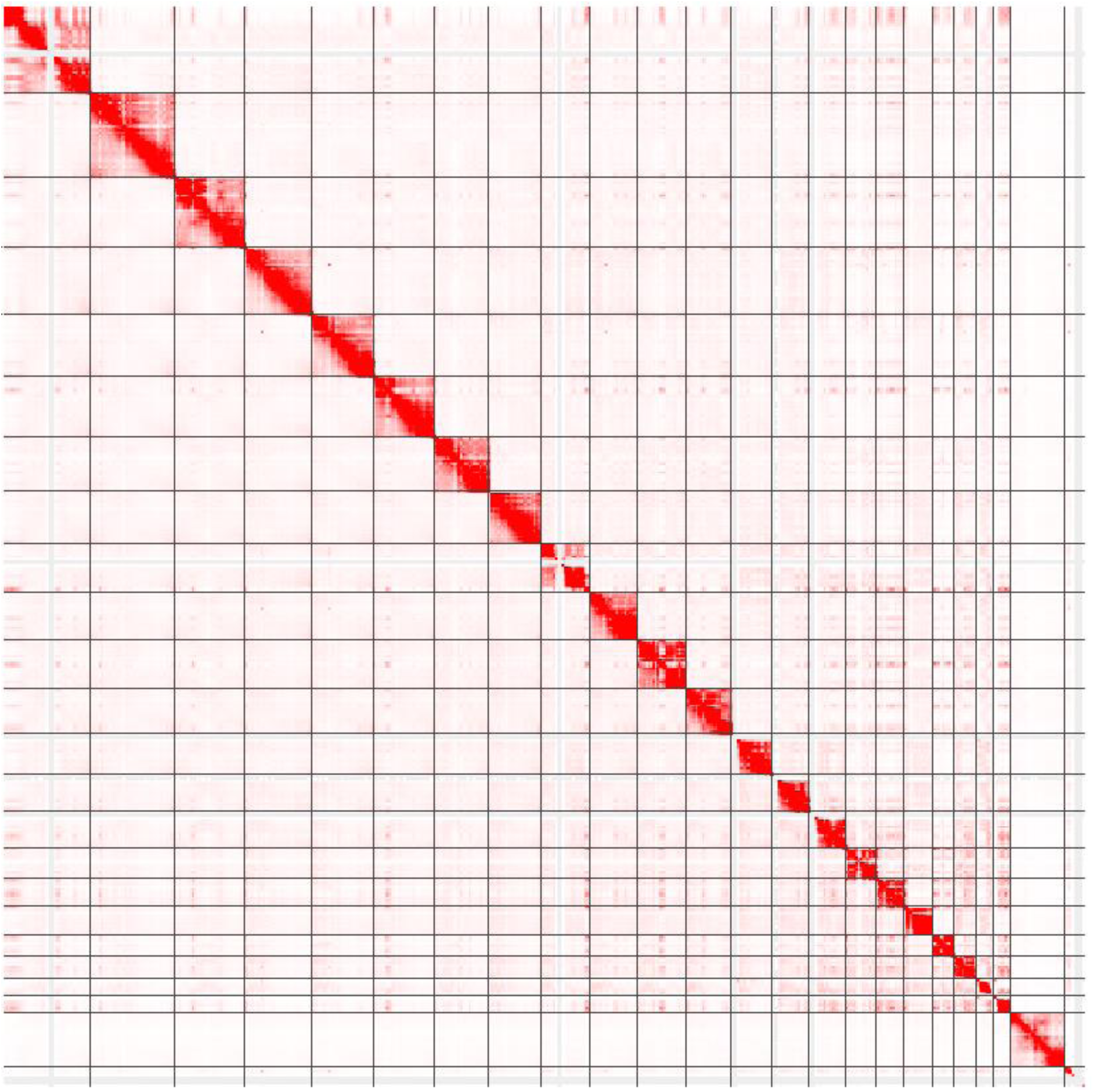
Hi-C data reveals regions of the genome in close proximity. Juicebox plot showing the contact frequency map (100kb resolution) for the entire genome with each chromosome laid end to end (X and Y chromosomes at the bottom right). Each red dot consists of two regions of the genome associated with each other. Top right: HC patient monocytes. Bottom Left: AH patient monocytes

Using HiCRep^23^, we measured the correlation coefficient of the chromatin interactions for each chromosome between every sample to assess in an unbiased manner how much variation there is in genome architecture between different patients. For most comparisons, the correlation coefficient between all individuals was very high (>90%), indicative of the high conservation of 3D genome architecture in monocytes regardless of disease status. But for most chromosomes, higher correlations were observed between healthy controls or between AH samples, and not between the two groups (**Figure 2**), indicative of a change in genome architecture caused by disease. Notable exceptions include chromosomes 4, 11, 16, 19 and the X chromosome. For the X chromosome, one female healthy control sample (HC2) differed significantly from the 7 males, and the 3 HC males were more similar to each other than the 4 AH patients (**Supplemental Figure 2**). Chromosomes 4, 11, and 19 mostly showed higher similarity by disease, except the female healthy control was more similar to the AH patients. Chromosome 16 did not segregate by disease. These data indicate that while much of the 3D genome architecture of monocytes does not change with disease, there likely exist chromosome specific regions that differ significantly.

**Figure 2:**
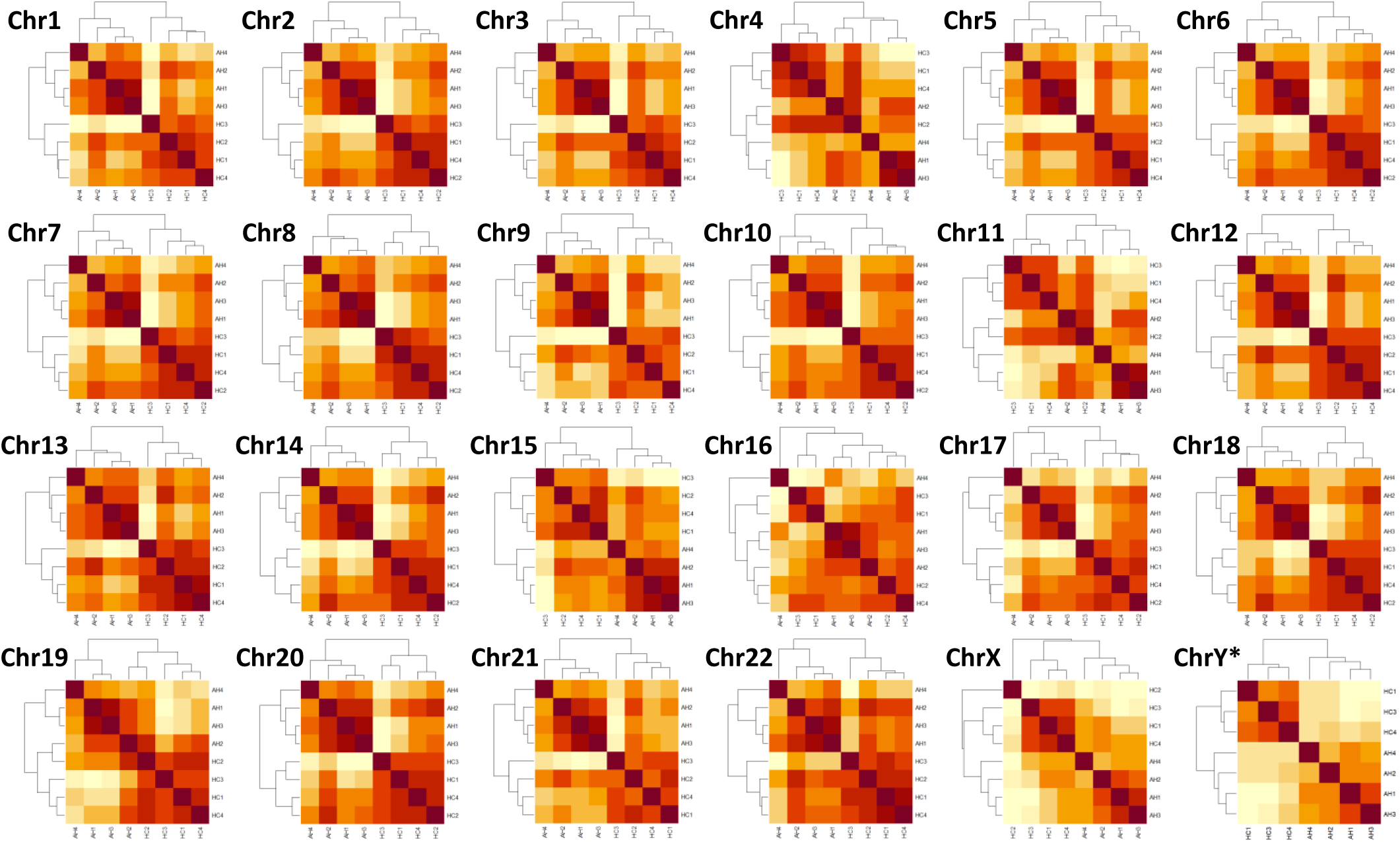
Correlation reveals differences between AH and HC. Hierarchically clustered heatmaps of correlation coefficients of the Hi-C data for each chromosome. For each heatmap, darker colors refer to higher correlation coefficients.

### Differences in 3D Architecture occur throughout all chromosomes in AH

Hi-C data reveal regions of the genome in close proximity via the contact frequency between two loci. We can measure disease-specific changes in 3D genome architecture by measuring statistically significant differences in these frequencies. Using multiHiCcompare^24^, we measured the contact frequency between every pair of 100kb genome windows (region-region pairs) and calculated statistically significant changes between HC and AH. We observed significant change in contact frequency throughout the entire length of each chromosome (**Figure 3**). To ensure the method was reliable, we randomized the samples 5 times prior to statistical testing and observed almost no significant changes between regions in any of the tests (**Supplemental Table 3**).

**Figure 3:**
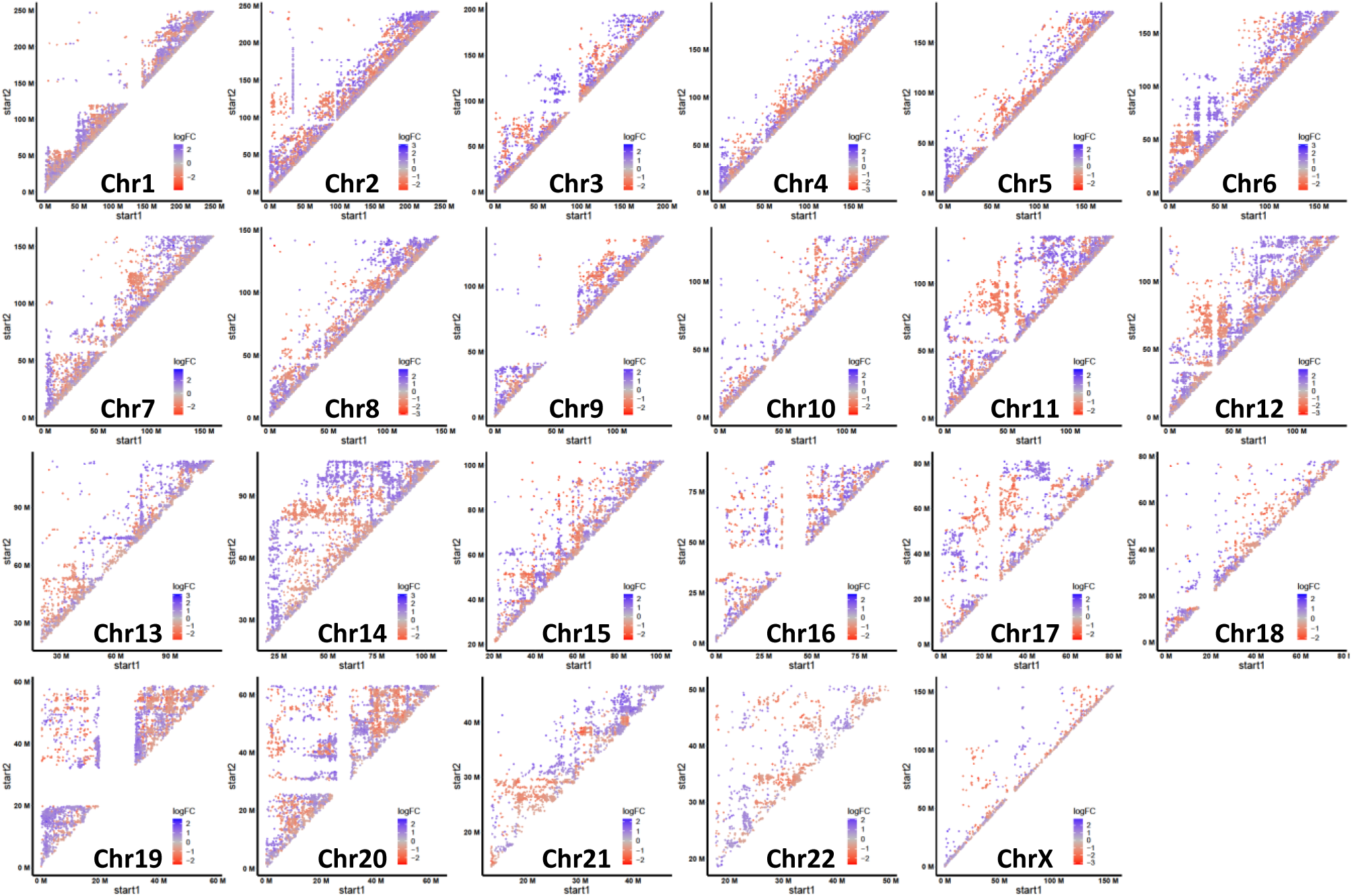
Changes in Contact Frequency in Disease are Local and Long-Range. Plots showing regions with significant changes in contact frequency in AH. Each dot indicates two regions of the genome that either increased (Blue) or decreased (Red) contact frequency in AH. X- and Y-axes are position along the chromosome

For most chromosomes, the changes in contact frequency caused by disease occurred in region-region pairs that are adjacent to each other, which corresponds to the fact that most Hi-C interactions are between local genomic regions (**Supplemental Figure 3**)^25,26^. For example, interaction data from Chromosome 2 is primarily between adjacent regions and diminishes by distance, and similarly changes in contact frequency caused by disease are also mostly local (**Supplemental Figure 4**). Chromosome 1 is an exception because it can be separated into 3 major megadomains, and changes in genome structure in chromosome 1 were restricted to these regions, with almost nothing differing between each megadomain (**Supplemental Figure 5**). Some chromosomes did have long-distance structural changes between chromosome arms, including chromosomes 11, 12, 14, 16, 17, 19, and 20. The telomeric regions of Chromosome 14 had extensive increases in contact frequency with the body of the chromosome in AH. The telomeres of Chromosome 14 contain the T-cell receptor alpha (TRA) locus on one arm and the immunoglobulin heavy chain (IGH) locus on the other.

### Numerous Hotspots with significant structural change in AH contain genes important for innate immunity

While there were differences in contact frequency throughout the genome in AH, there were a number of notable “hotspots” that contained a high density of the most significantly altered regions (**Figure 4A**). To identify these hotspots, we focused only on regions of the genome with exceptionally high statistically significant changes in contact frequency (padj<0.0000001). From this list of we identified 22 “hotspots” regions of the genome with a high frequency of these changes caused by disease (**Table 2, Supplemental Table 4**). The size of these hotspots varied significantly from 21MB down to less than 1MB. For most hotspots, the differences in contact frequency occurred entirely within the region. Many of the hotspots entirely increased or decreased contact frequency in AH while some contained separate areas that increased or decreased, further indicative of a localized rearrangement in genome structure, with some areas moving closer and others moving further away in disease.

**Figure 4:**
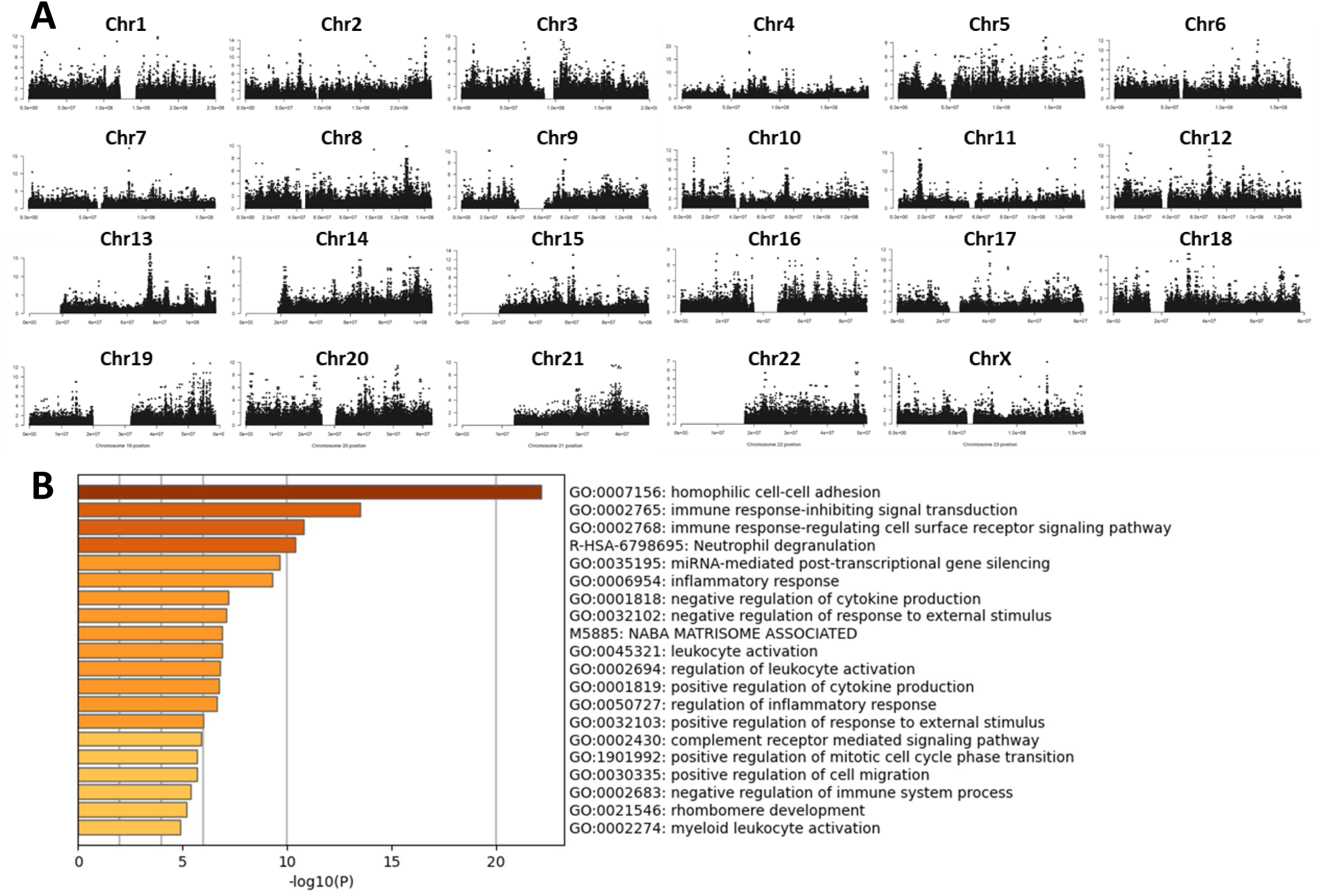
Certain Regions in the Genome have Hotspots of Significant Change in Disease. A)Manhattan plot showing regions of the genome with significant changes in chromatin interaction with disease. X-axis is position along the chromosome. Y-axis is –log10(p). B) Pathway analysis of genes in regions with the most significant changes with disease.

**Table 2:**
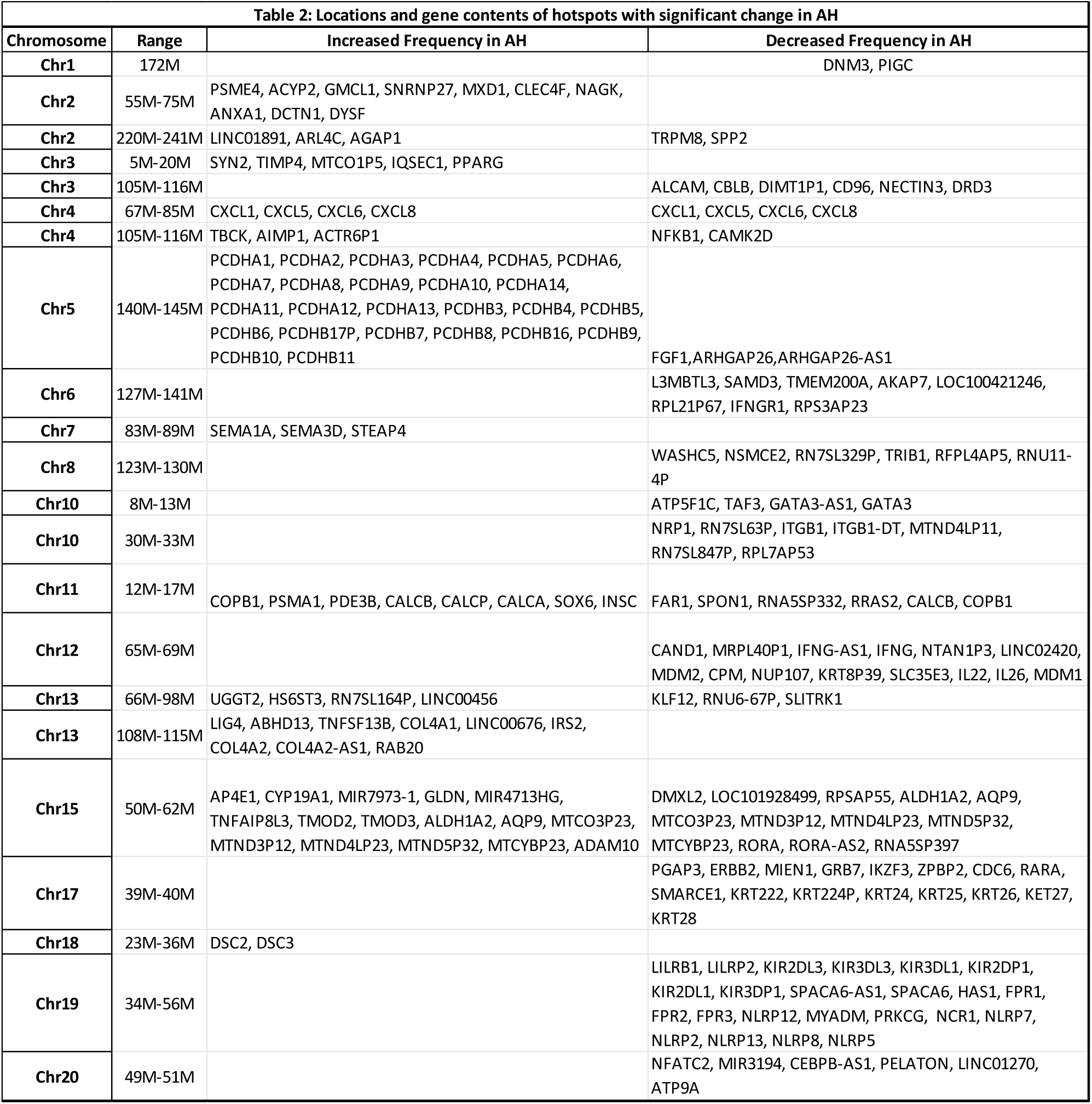
Locations and gene contents of hotspots with significant change in AH.

Considering their size, some hotspots contained hundreds of genes, including many genes associated with immune responses and have been associated with AH (**Table 2, Supplemental Tables 5 and 6**). We performed pathway analysis on all of the genes in the hotspots and in regions with very significant changes in structure (all genes in **Supplemental Table 4**) and found that these regions are enriched for innate immune signaling (**Figure 4B**). Many of the hotspots contain large families of genes that are clustered together in the genome. For example, Chromosome 4 (∼67-85M) contains the CXC-chemokines, Chromosome 5 (∼140-145M) contains the protocadherins, and Chromosome 12 (∼65-69M) contains IFNG, IL-22, and IL-26. One hotspot on Chromosome 19 (∼34-56M) contains multiple large families of genes that are involved in diverse aspects of immune regulation including killer-cell immunoglobulin-like receptors (KIRs), leukocyte immunoglobulin-like receptors (LILRs), sialic-acid-binding immunoglobulin-like lectins (Siglecs), C5 complement receptors, and NOD-like family of receptors (NLRPs). Additionally, hotspots also contained other notable immune genes like NFKB1 (Chromosome 4), IFNGR1 (Chromosome 6), Integrin Beta (ITGB1) (Chromosome 1), and NFATC2 (Chromosome 20).

### Structural Changes Observed at Innate Immune Gene Clusters are more complex than TADs

Because hotspots represent regions of the genome that have a large number of structural changes in disease, and they contain important genes involved in innate immunity, these regions were examined with finer detail. Within the hotspots, TAD structure, such as the loss or expansion of a TAD, was unchanged. Instead, most changes in genome architecture were due to the interaction profile within individual TADs and the relationship between the TADs and the surrounding genome.

CXC-chemokines are upregulated in AH, and we previously found increased co-regulation of these genes in response to LPS in AH patients, using single cell data of monocytes^20^. Zooming into the hotspot on chromosome 4 containing the CXC-chemokine cassette, we see that all of the CXC-chemokine genes exist within a single TAD (**Figure 5**). Interestingly, while the borders of the TAD did not change, the contact frequency within that TAD increased in AH. Zooming out, we observed an alternating pattern of interactions between this TAD and the neighboring TADs.

**Figure 5:**
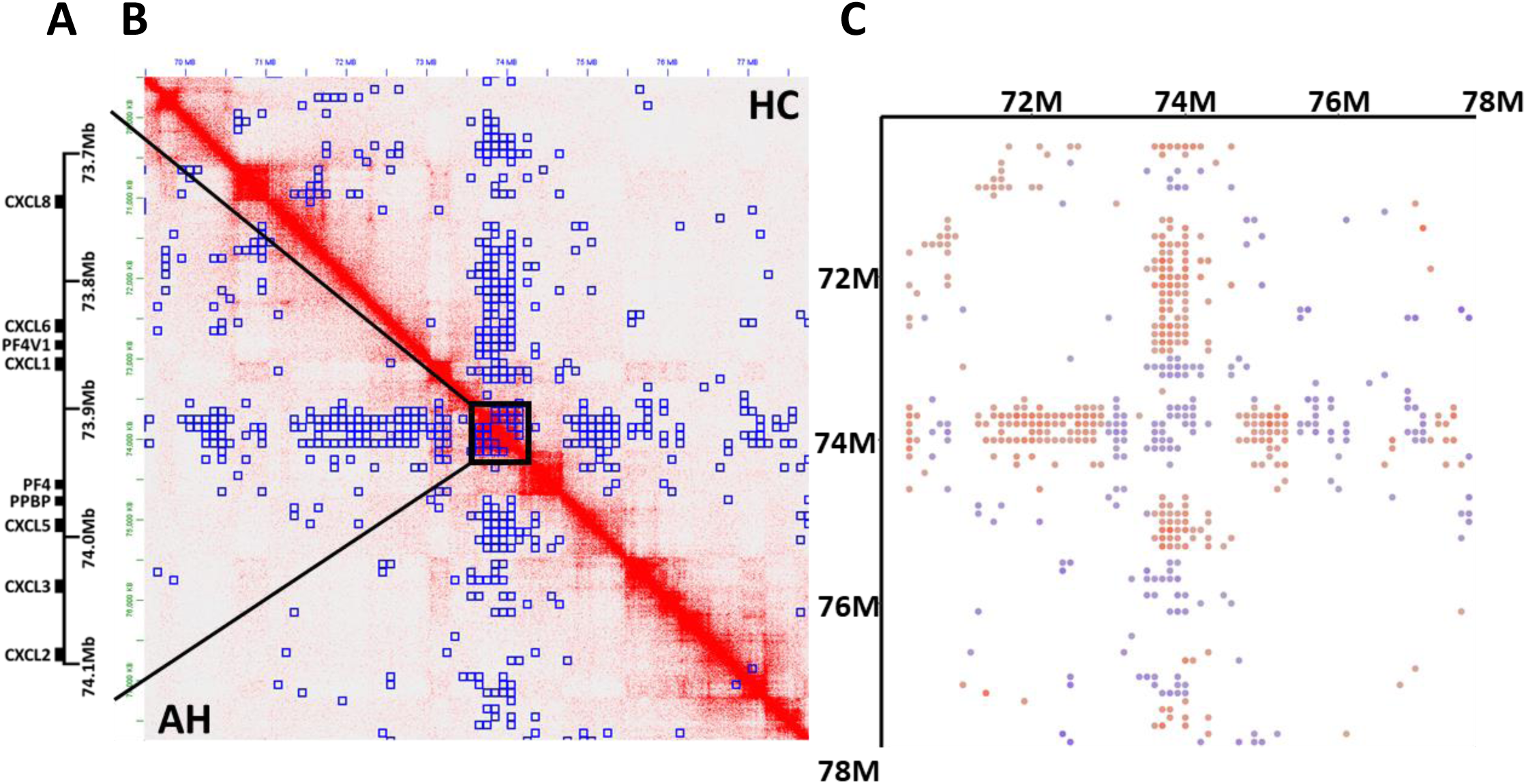
CXC-chemokine cluster has increase contact frequency within the TAD and reduced outside. **A)** Schematic of the CXC-chemokine gene cluster. **B)** Juicebox plot showing contact frequency map of the CXC-chemokine gene cluster and the surrounding genomic region. Blue boxes are 100kb regions significantly different in disease. **C)** Plot corresponding to the Juicebox plot showing whether each significant region increased (Blue) or decreased (Red) contact frequency in disease.

We previously found that monocytes in AH have increased expression of CLRs, which are a family of PRRs that sense a wide diversity of PAMPs and DAMPs. Many of these CLRs are found within the NK gene receptor complex on chromosome 12, and in particular, four of these CLRs (Mincle, Dectin-2, Dectin-3 and DCIR) are in close proximity and have highly co-regulated expression in AH. Similar to the CXC-chemokines, TAD structure in this hotspot did not change, but rather there were increased interactions within the individual TADs, increased connections between TADs within the entire NK-gene receptor complex (∼7M-10M), and reduced interaction immediately outside of it (**Figure 6**). Another hotspot on chromosome 12 at approximately 65-69M, which contains the genes for IL-22, IL-26, and IFNG, had the opposite structure, with fewer connections within the hotspot and more connections to the surrounding genomic landscape (**Figure 7**).

**Figure 6:**
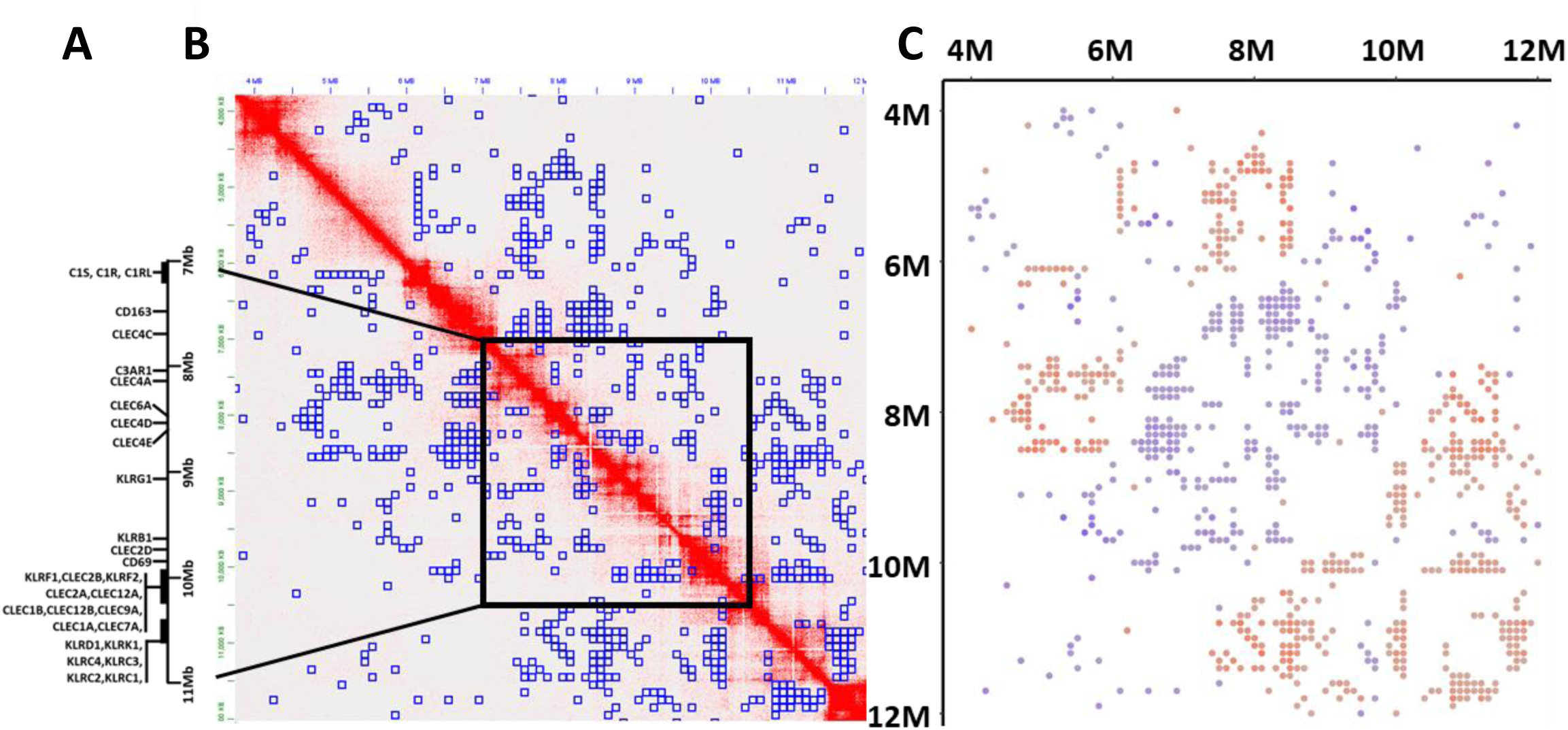
Myeloid CLR gene cluster has increased local contacts in AH. **A)** Schematic of the NK-gene receptor complex. **B)** Juicebox plot showing contact frequency map of the NK-gene receptor complex and the surrounding genomic region. Blue boxes are 100kb regions significantly different in disease. **C)** Plot corresponding to the Juicebox plot showing whether each significant region increased (Blue) or decreased (Red) contact frequency in disease.

**Figure 7:**
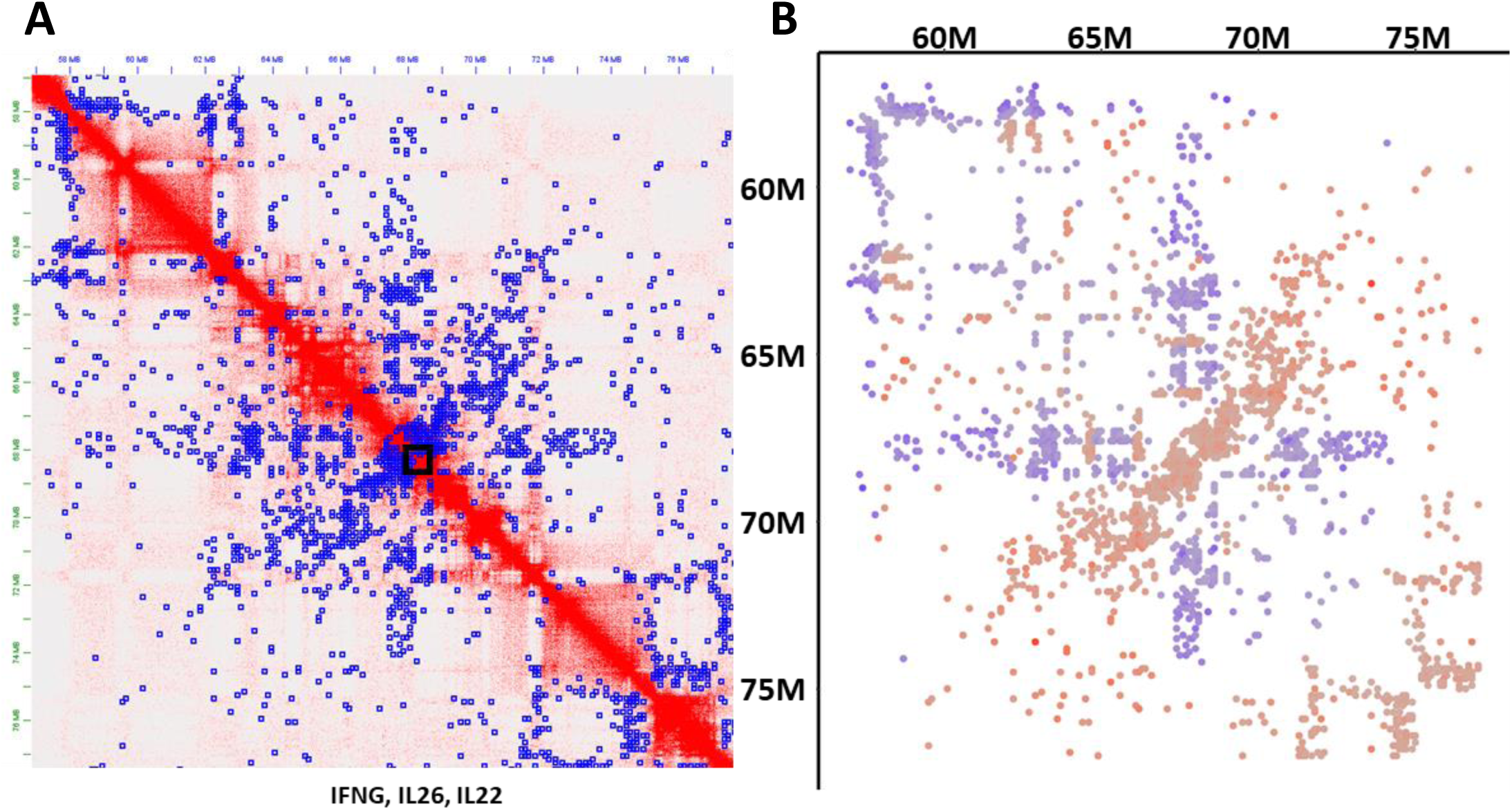
Genomic region around IL-22 and IFNG have decreased local contacts in AH. **A)** Juicebox plot showing contact frequency map of the region on Chromosome 12 around the IL-22 and IFNG genes. Blue boxes are regions significantly different in disease. **B)** Plot corresponding to the Juicebox plot showing whether each significant region increased (Blue) or decreased (Red) contact frequency in disease.

### Two adjacent loci of CCL chemokine genes have increased connectivity and correlated expression

In AH, CC-chemokines are upregulated in monocytes, similar to CXC-chemokines and many other pro-inflammatory cytokines^20,27^. In the genome, the CC-chemokine genes are in two loci separated by about 1.5MB. From the Hi-C data, each CC-chemokine locus is within an individual TAD (**Figure 8**). While this region of the genome was not a hotspot, there was significant changes in 3D genome architecture, particularly in terms of the relationship between the CC-chemokine loci and the surrounding area, which increased connectivity in disease.

**Figure 8:**
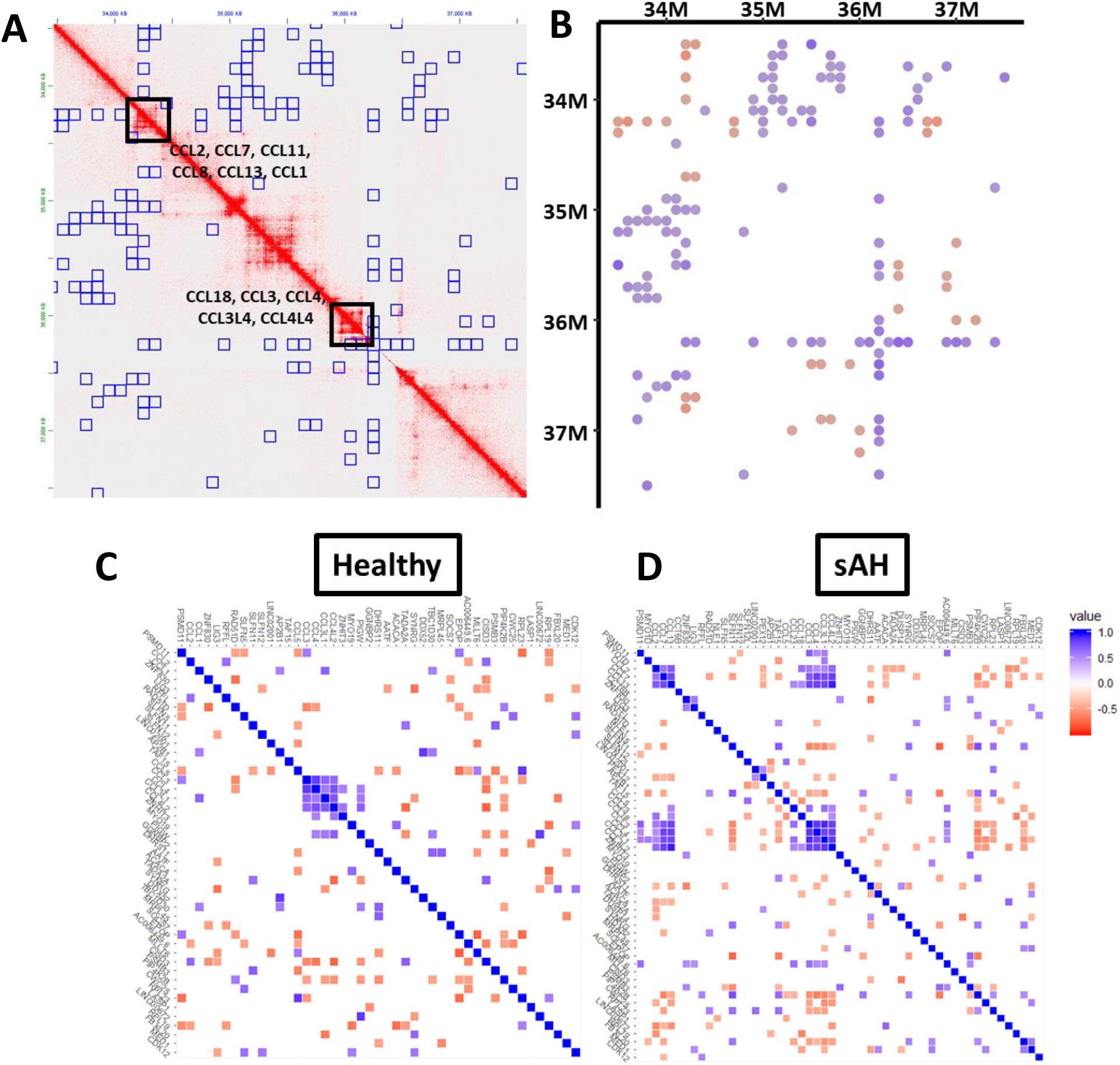
The Split CC-Chemokine Cluster has increased contact between loci, and highly correlated expression in AH. **A)** Juicebox plot showing contact frequency map of the two CC-chemokine gene clusters and the surrounding genomic region. Blue boxes are regions significantly different in disease. **B)** Plot corresponding to the Juicebox plot showing significantly increased (Blue) or decreased (Red) contact frequency in disease. **C/D)** Correlation analysis showing coordinately expressed in response to LPS (100pg/ml, for 24 hours) for healthy control (C) and AH (D). Genes are organized by chromosomal position and oriented in the same manner as A and B. Blue squares indicate genes with highly correlated expression while red squares indicate anti-correlated expression.

We previously published single-cell RNA-seq data from PBMCs isolated from patients with severe AH and healthy controls, with and without *ex vivo* LPS challenge (100pg/ml for 24 hours). In response to LPS, both HC and AH monocytes express chemokines, though the expression is much higher in AH. Next, we measured how well correlated gene expression was in single cells, in order to determine if neighboring genes have coordinated expression (**Figure 8**). In HC, we observed highly co-regulated expression of the CCL-chemokines in one of the loci (CCL3, CCL4, CCL3L1, CCL4L2). In AH, CCL-chemokine expression was highly co-regulated in both loci, including a few more genes in the first locus (CCL23, CCL18, CCL3, CCL4, CCL3L1, CCL4L2) and genes in the second locus (CCL2, CCL7, CCL13). And importantly, expression between both loci were also coordinated, suggesting the change in 3D genome architecture allowed for coordinated expression of these genes.

## Discussion

Alcohol-associated Hepatitis is characterized by dysfunctional monocytes that increase systemic inflammation and can cause extensive damage to the liver and eventually end-organ failure. Monocytes from AH patients are hypersensitive to innate immune stimuli, and in response to bacterial LPS, upregulate expression of cytokines and chemokines much more dramatically than seen in healthy controls. In this study, we try to understand how changes in the 3D genome architecture of monocytes are transformed during AH and infer the impact these changes have on gene expression.

Many studies have been conducted trying to understand how changes in 3D genome architecture affects gene expression and cell development using unbiased and untargeted methods like Hi-C. Far fewer studies have been conducted on primary monocytes and how the architecture changes with disease. We hypothesized that in a disease like AH, where immune cells are so hypersensitive to LPS, alterations in 3D genome architecture may play a significant role in gene regulation. In our study, using monocytes isolated from four AH patients and 4 healthy controls, we in fact see significant changes in chromatin conformation caused by disease. But by measuring significant changes in contact frequency for every region-region pair, we were able to assess that throughout each chromosome, there were significant changes in how the TADs and larger loci associated with each other. Notably, we found a number of hotspots in the genome that contained a large number of structural changes.

Many large gene families and genes associated with innate immunity and AH were found within these hotspots. For example, in a previous publication, our group found highly coordinated expression of CXC-chemokines and some of the C-type lectin genes present within the NK-gene receptor complex^20^. Here, we find that the 3D genome architecture of these regions has significantly changed in disease. Moreover, throughout these hotspots we identified a number of other genes and gene families associated with AH. Chromosome 19 contained a large hotspot containing genes associated with the inflammasome (NOD-like receptors), which is an important inflammatory complex involved in IL-1B release and pyroptosis, which contributes to significant damage in liver and other organs during AH^28,29^. This hotspot also contained SPACA6, which is a host gene for a cluster of microRNAs that are also upregulated in AH^30^.

This region on chromosome 19 also contains the killer cell immunoglobulin-like receptor family, which are genes that encode transmembrane glycoproteins involved in NK-cell target recognition. Alongside the NK-gene receptor complex on Chromosome 12, another hotspot, our data suggests significant architectural changes in two different loci important for NK-cell target recognition in monocytes. While NK-cells are dysfunctional in AH^31,32^, the structural changes we observe in monocytes in these regions is more likely to do with the other gene families in the area, such as in the chromosome 12 hotspot, the CLR genes, and in the chromosome 19 hotspot, the leukocyte immunoglobulin-like receptors, which are highly coordinately expressed in monocytes.

On the other hand, some of the hotspots contained genes with unclear roles in monocytes or AH. Looking at single cell data, some of these hotspots contained genes with very low expression in monocytes. For example, the T-cell receptor locus on chromosome 14, which increased contact frequency throughout the length of the chromosome, but none of these genes are expressed in monocytes. But all of these hotspots are more than just the genes within and may contain important regulatory elements.

While the CC-chemokine gene cassettes on chromosome were not a clear hotspot with a large number of changes in disease, there were still significant differences. This region was of interest because many CCL chemokines are highly upregulated in response to LPS in AH patients^20^, and levels of circulating CCL chemokines are higher in disease^33^. In AH, the two CC-chemokine cassettes, which are separated by a little more than 1Mb and are in different TADs, had higher contact frequency and from single-cell data, had highly correlated expression. This suggests that these two regions came closer together in disease, and that the proximity of regulatory elements had an effect on the expression of these genes.

While there were significant changes in 3D genome architecture in AH, this study has a number of limitations. Other Hi-C studies in monocytes and THP-1 cells have been limited by small sample size, typically one patient/sample or two patients with the data combined to increase resolution. This is not ideal for human disease which is typically multifactorial and complex. In this study, we studied monocytes isolated from four healthy controls and four AH patients, in order to look at changes in contact frequency with high statistical confidence. But the trade-off was that we could not analyze this data at a much higher resolution without much deeper sequencing. Still, to fully understand how genome architecture changes with disease, even more patient samples need to be sequenced and analyzed with Hi-C, to encompass the multifactorial nature of this complex human disease.

Taken together, these results indicate that the 3D genome architecture of monocytes is significantly altered in AH and suggest that perturbations in the 3D architecture contribute to differences in gene expression, especially in response to innate immune challenges. Future studies will focus on trying to understand the cause of genome restructuring in AH, as both alcohol and innate immune stimuli have the potential to alter epigenetic gene regulation^34^. Additionally, more work is needed to understand how changes in local genome structure affect transcription factor dynamics and the relationship of enhancers and promoters in disease. Finally, these studies suggest that Hi-C is a useful tool to understand how disease alters the genome, and there is an ongoing need to build more databases of this kind of data from a wider diversity of cell types, disease states, and patients.

## Methods

### Alcohol-related Hepatitis and Healthy Control Patient Selection

Enrolled patients had confirmed diagnosis of AH by clinicians at the Cleveland Clinic based on medical history, physical examination, and laboratory results, according to the guidelines of the American College of Gastroenterology [https://gi.org/clinical-guidelines/] (**Supplemental Table 1**). Healthy controls were recruited from the Clinical Research Unit at the Cleveland Clinic.

### Isolation of Human PBMCs

PBMCs were isolated from human blood as previously described^20,30^. Isolation of mononuclear cells was performed by density gradient centrifugation on Ficoll-Paque PLUS (GE Healthcare, Uppsala, Sweden). 1 mL of freshly collected Buffy Coat was mixed at a ratio of 1:5 (vol/vol) with phosphate buffered saline (PBS) at 37°C and divided and layered onto 8 mL Ficoll-Paque PLUS in two 15 mL conical centrifuge tubes. After centrifugation at 400 × g for 30 min at 20°C (no brake), buffy coat fractions were collected, pooled, resuspended in culture media (Roswell Park Memorial Institute (RPMI)-1640 supplemented with 100 μM Penicillin-streptomycin and 10% fetal bovine serum (FBS)), and centrifuged at 400 × g for 15 min at 20°C. The pellets were resuspended in 8 mL of culture media, counted, and again centrifuged at 400 × g for 8 min at 20°C. Cells were then cryo-preserved by resuspending in freezing media (50% culture media, 40% FBS, 10% dimethyl sulfoxide (DMSO)) at a concentration of 1.5 × 10^6^ cells/mL, and allowed to freeze slowly to −80°C in a styrofoam container. For long-term preservation, cells were stored in liquid nitrogen.

### Hi-C of Isolated Monocytes from Cryopreserved PBMCs

Cryopreserved PBMCs from 4 AH patients and 4 age-matched healthy controls were thawed following the 10x protocol for cryopreserved PBMCs. Monocytes were isolated by negative selection using the EasySep Human Monocyte Enrichment Kit without CD16 Depletion (StemCell Technologies, Cambridge, MA) according to factory instructions. Hi-C was performed with the Arima Genome-wide Hi-C kit (Carlsbad, CA) according to factory instructions. Libraries were pooled and sequenced with an Illumina Novaseq 6000.

### Analysis of Hi-C Data

Sequencing data for the Hi-C experiments were aligned to the genome (GRC38, release 93) using Juicer^35^. From the output of Juicer, we summarized the quality of the Hi-C maps using the inter.txt files (**Supplementary Table 2**). Chromosomal correlations were calculated using HiCRep^23^. Differential contact frequencies were measured using multiHiCcompare, at a resolution of 100kb and using the inter.hic files from Juicer^24^. Hi-C data was visualized using Juicebox to view each of the inter.hic files^36^. Regions with changes in contact frequency in disease were labeled using data from multiHiCcompare. To ensure these changes in contact frequency are due to disease and not random chance, we randomized the samples and recalculated differential contact frequency and found significantly fewer differences (**Supplemental Table 3**).

“Hotspots” were defined as regions of the genome with a high frequency of changes in contact frequency. To identify these regions, we took the list of all differential contacts from multiHiCcompare and filtered by a very stringent statistical cutoff of padj<0.0000001. Then we identified regions of the genome with a large number of these differential contacts, which identified 22 “hotspots” (**Table 2, Supplemental Table 4**). To identify the borders of these regions, we looked more liberally at the full list of differential contacts (padj<0.05). Gene enrichment for genes present in hotspots and other highly differential regions was performed using Metascape^37^.

### scRNA-seq Analysis and Clustering

Sequencing data was aligned to the Human genome (GRC38, release 93) using cellranger (v3.0.2). All gene expression and clustering analyses were performed using Seurat (3.1.1) as previously described^20,38^. Briefly, all samples were first normalized using SCTransform and then filtered to remove low quality cells (nFeature_RNA<4000, nFeature_RNA>200, percent.mt<20, which removes doublets, cells with low reads, and cells with high mitochondrial content))^39^. All samples were combined using the PrepSCTIntegration and FindIntegrationAnchors functions to find common anchor genes in all samples for all cell types, and then integrated using the IntegrateData function, with all normalizations using the SCT transformed data^38,40^. Clustering was performed using RunPCA and RunUMAP, and clusters were identified using FindNeighbors and FindClusters.

### bigSCale2

For correlation analyses, bigSCale2 was used to calculate correlations using the Z-score algorithm^41^. For these analyses, only monocytes were used (clusters labeled CD14_Monocyte1, CD14_Monocyte2, CD14_Monocyte3, and CD16_Monocyte). All gene clusters were determined by first ordering all annotated human genes by chromosome and start codon then finding the desired clusters and selecting an arbitrary set of genes surrounding it (genes that are not hypothesized to be involved) to ensure the entire cluster was obtained. Genes were then filtered for low expression using the criteria that bigSCale2 was unable to determine a correlation coefficient. Heatmaps of the correlation coefficients were made with only the top and bottom 5% of all correlation coefficients shown, to remove noise and isolate the most important correlations.

## Data and Code Availability

The Hi-C data generated for this study can be found at the database of Genotypes and Phenotypes (dbGaP) [TBD] The scRNA-seq data for this study can be found at National Center for Biotechnology Information Gene Expression Omnibus under accession number [PRJNA596980]^20^. All scripts used for analyses, differential expression results, for all cell types, and figure generation can be found at the author’s github (https://github.com/atomadam2/). Any additional information required to reanalyze the data reported in this paper is available from the lead contact upon request.

## Ethical Approval and Consent to Participate

The study protocol was approved by the Institutional Review Board for the Protection of Human Subjects in Research at the Cleveland Clinic and MetroHealth Hospitals, Cleveland. All methods were performed in accordance with the IRB’s guidelines and regulations and written informed consent was obtained from all subjects.

## Author Contributions

AK contributed conception and design of the study. MRM helped conduct the experiments. JD, AB, DS, NW, SD recruited patients and performed clinical analyses. AK analyzed the data and wrote the first draft of the manuscript. All authors contributed to manuscript revision, read and approved the submitted version.

## Acknowledgements

The authors thank the Clinical Research Unit, Genomics Core and Computing Services at the Cleveland Clinic. Additionally, Dr. Laura E Nagy, from the Cleveland Clinic and Northern Ohio Alcohol Center, for reagents and critical comments on the manuscript.

## References

1 Cremer, T. & Cremer, C. Chromosome territories, nuclear architecture and gene regulation in mammalian cells. Nat Rev Genet 2, 292–301 (2001). 10.1038/35066075

2 Szabo, Q., Bantignies, F. & Cavalli, G. Principles of genome folding into topologically associating domains. Sci Adv 5, eaaw1668 (2019). 10.1126/sciadv.aaw1668

3 Sanborn, A. L. et al. Chromatin extrusion explains key features of loop and domain formation in wild-type and engineered genomes. Proc Natl Acad Sci U S A 112, E6456–6465 (2015). 10.1073/pnas.1518552112

4 Vietri Rudan, M., et al. Comparative Hi-C reveals that CTCF underlies evolution of chromosomal domain architecture. Cell Rep 10, 1297–1309 (2015). 10.1016/j.celrep.2015.02.004

5 Rao, S. S. P. et al. Cohesin Loss Eliminates All Loop Domains. Cell 171, 305–320 e324 (2017). 10.1016/j.cell.2017.09.026

6 Nora, E. P. et al. Targeted Degradation of CTCF Decouples Local Insulation of Chromosome Domains from Genomic Compartmentalization. Cell 169, 930–944 e922 (2017). 10.1016/j.cell.2017.05.004

7 Schwarzer, W. et al. Two independent modes of chromatin organization revealed by cohesin removal. Nature 551, 51–56 (2017). 10.1038/nature24281

8 Stik, G. et al. CTCF is dispensable for immune cell transdifferentiation but facilitates an acute inflammatory response. Nat Genet 52, 655–661 (2020). 10.1038/s41588-020-0643-0

9 Yang, B. et al. CTCF controls three-dimensional enhancer network underlying the inflammatory response of bone marrow-derived dendritic cells. Nat Commun 14, 1277 (2023). 10.1038/s41467-023-36948-5

10 Kolovos, P. et al. Binding of nuclear factor kappaB to noncanonical consensus sites reveals its multimodal role during the early inflammatory response. Genome Res 26, 1478–1489 (2016). 10.1101/gr.210005.116

11 Liu, R. et al. Hi-C, a chromatin 3D structure technique advancing the functional genomics of immune cells. Front Genet 15, 1377238 (2024). 10.3389/fgene.2024.1377238

12 Cuartero, S., Stik, G. & Stadhouders, R. Three-dimensional genome organization in immune cell fate and function. Nat Rev Immunol 23, 206–221 (2023). 10.1038/s41577-022-00774-5

13 Platanitis, E. et al. Interferons reshape the 3D conformation and accessibility of macrophage chromatin. iScience 25, 103840 (2022). 10.1016/j.isci.2022.103840

14 Minderjahn, J. et al. Postmitotic differentiation of human monocytes requires cohesin-structured chromatin. Nat Commun 13, 4301 (2022). 10.1038/s41467-022-31892-2

15 Xia, Y. et al. Capturing 3D Chromatin Maps of Human Primary Monocytes: Insights From High-Resolution Hi-C. Front Immunol 13, 837336 (2022). 10.3389/fimmu.2022.837336

16 Zhang, Z. et al. Massive reorganization of the genome during primary monocyte differentiation into macrophage. Acta Biochim Biophys Sin (Shanghai) 52, 546–553 (2020). 10.1093/abbs/gmaa026

17 Liu, Y., Li, H., Czajkowsky, D. M. & Shao, Z. Monocytic THP-1 cells diverge significantly from their primary counterparts: a comparative examination of the chromosomal conformations and transcriptomes. Hereditas 158, 43 (2021). 10.1186/s41065-021-00205-w

18 Wang, M. et al. Chronic alcohol ingestion modulates hepatic macrophage populations and functions in mice. J Leukoc Biol 96, 657–665 (2014). 10.1189/jlb.6A0114-004RR

19 Ju, C. & Mandrekar, P. Macrophages and Alcohol-Related Liver Inflammation. Alcohol Res 37, 251–262 (2015).

20 Kim, A., Bellar, A., McMullen, M. R., Li, X. & Nagy, L. E. Functionally Diverse Inflammatory Responses in Peripheral and Liver Monocytes in Alcohol-Associated Hepatitis. Hepatol Commun 4, 1459–1476 (2020). 10.1002/hep4.1563

21 Wu, X. et al. Recent Advances in Understanding of Pathogenesis of Alcohol-Associated Liver Disease. Annu Rev Pathol 18, 411–438 (2023). 10.1146/annurev-pathmechdis-031521-030435

22 Liu, M. et al. Super enhancer regulation of cytokine-induced chemokine production in alcoholic hepatitis. Nat Commun 12, 4560 (2021). 10.1038/s41467-021-24843-w

23 Yang, T. et al. HiCRep: assessing the reproducibility of Hi-C data using a stratum-adjusted correlation coefficient. Genome Res 27, 1939–1949 (2017). 10.1101/gr.220640.117

24 Stansfield, J. C., Cresswell, K. G. & Dozmorov, M. G. multiHiCcompare: joint normalization and comparative analysis of complex Hi-C experiments. Bioinformatics 35, 2916–2923 (2019). 10.1093/bioinformatics/btz048

25 Burton, J. N. et al. Chromosome-scale scaffolding of de novo genome assemblies based on chromatin interactions. Nat Biotechnol 31, 1119–1125 (2013). 10.1038/nbt.2727

26 Lieberman-Aiden, E. et al. Comprehensive mapping of long-range interactions reveals folding principles of the human genome. Science 326, 289–293 (2009). 10.1126/science.1181369

27 Degre, D. et al. Hepatic expression of CCL2 in alcoholic liver disease is associated with disease severity and neutrophil infiltrates. Clin Exp Immunol 169, 302–310 (2012). 10.1111/j.1365-2249.2012.04609.x

28 Khanova, E. et al. Pyroptosis by caspase11/4-gasdermin-D pathway in alcoholic hepatitis in mice and patients. Hepatology 67, 1737–1753 (2018). 10.1002/hep.29645

29 Petrasek, J. et al. IL-1 receptor antagonist ameliorates inflammasome-dependent alcoholic steatohepatitis in mice. J Clin Invest 122, 3476–3489 (2012). 10.1172/JCI60777

30 Kim, A., Saikia, P. & Nagy, L. E. miRNAs Involved in M1/M2 Hyperpolarization Are Clustered and Coordinately Expressed in Alcoholic Hepatitis. Frontiers in Immunology 10 (2019). 10.3389/fimmu.2019.01295

31 Kim, A. et al. Diminished function of cytotoxic T- and NK-cells in severe alcohol-associated hepatitis. Metabolism and Target Organ Damage 2, 18 (2022). 10.20517/mtod.2022.13

32 Stoy, S. et al. Cytotoxic T lymphocytes and natural killer cells display impaired cytotoxic functions and reduced activation in patients with alcoholic hepatitis. Am J Physiol Gastrointest Liver Physiol 308, G269–276 (2015). 10.1152/ajpgi.00200.2014

33 Luther, J., Vannier, A. G., Schaefer, E. A. & Goodman, R. P. The circulating proteomic signature of alcohol-associated liver disease. JCI Insight 7 (2022). 10.1172/jci.insight.159775

34 Malherbe, D. C. & Messaoudi, I. Transcriptional and Epigenetic Regulation of Monocyte and Macrophage Dysfunction by Chronic Alcohol Consumption. Front Immunol 13, 911951 (2022). 10.3389/fimmu.2022.911951

35 Durand, N. C. et al. Juicer Provides a One-Click System for Analyzing Loop-Resolution Hi-C Experiments. Cell Syst 3, 95–98 (2016). 10.1016/j.cels.2016.07.002

36 Robinson, J. T. et al. Juicebox.js Provides a Cloud-Based Visualization System for Hi-C Data. Cell Syst 6, 256–258 e251 (2018). 10.1016/j.cels.2018.01.001

37 Zhou, Y. et al. Metascape provides a biologist-oriented resource for the analysis of systems-level datasets. Nat Commun 10, 1523 (2019). 10.1038/s41467-019-09234-6

38 Stuart, T. et al. Comprehensive Integration of Single-Cell Data. Cell 177, 1888–1902 e1821 (2019). 10.1016/j.cell.2019.05.031

39 Hafemeister, C. & Satija, R. Normalization and variance stabilization of single-cell RNA-seq data using regularized negative binomial regression. Genome Biol 20, 296 (2019). 10.1186/s13059-019-1874-1

40 Ding, J. et al. Systematic comparative analysis of single cell RNA-sequencing methods. bioRxiv, 632216 (2019). 10.1101/632216

41 Iacono, G., Massoni-Badosa, R. & Heyn, H. Single-cell transcriptomics unveils gene regulatory network plasticity. Genome Biol 20, 110 (2019). 10.1186/s13059-019-1713-4

